# The global proteome and phosphoproteome landscape of sepsis-induced kidney injury

**DOI:** 10.1101/2020.05.21.108464

**Authors:** Yi-Han Lin, Maryann P. Platt, Norberto Gonzalez-Juarbe, Dong Zhou, Yanbao Yu

**Author notes:** Equal contribution (Norberto Gonzalez-Juarbe, Dong Zhou, Yanbao Yu). Correspondance: Yanbao Yu,; Dong Zhou,; Norberto Gonzalez-Juarbe.

## Abstract

Sepsis-induced acute kidney injury (S-AKI) is the most common complication in hospitalized and critically ill patients, highlighted by a rapid decline of kidney function occurring a few hours or days after sepsis onset. Systemic inflammation elicited by microbial infections is believed to lead to kidney damage under immunocompromised conditions. However, while AKI has been recognized as a disease with long-term sequelae, partly due to the associated higher risk of chronic kidney disease (CKD), the understanding of kidney pathophysiology at the molecular level and the global view of dynamic regulations *in situ* after S-AKI, including transition to CKD, remains limited. Existing studies of S-AKI mainly focus on deriving sepsis biomarkers from body fluids. In the present study, we constructed a mid-severity septic murine model using cecal ligation and puncture (CLP), and examined the temporal changes to the kidney proteome and phosphoproteome at day 2 and day 7 after CLP surgery, corresponding to S-AKI and the transition to CKD, respectively by employing an ultrafast and economical filter-based sample processing method combined with the label-free quantitation approach. Collectively, we identified 2,119 proteins and 2,950 phosphosites through multi-proteomics analyses. Here we denote the pathways that are specifically responsive to S-AKI and its transition to CKD, which include regulation of cell metabolism regulation, oxidative stress, and energy consumption in the diseased kidneys. Our data can serve as an enriched resource for the identification of mechanisms and biomarkers for sepsis-induced kidney diseases.

## Introduction

Acute kidney injury (AKI) is characterized by a rapid decline of renal function resulting in elevated serum creatinine levels with or without decreased urine output ^1^. It is estimated that the incidence of AKI is about 10-15% of all patients admitted to the hospital, and its frequent occurrence with other organ dysfunction together causes high mortality ^2^. AKI has been recognized as a disease with long-term sequelae because survivors of AKI have a high risk of developing chronic kidney disease (CKD), which constitutes another high healthcare burden worldwide ^3^. In the clinic, the leading cause of AKI is sepsis, which is a systemic inflammatory response elicited by microbial infections ^4^. In the intensive care unit (ICU), sepsis accounts for up to 50% of AKI cases ^5^, and sepsis-induced AKI (S-AKI) has 6-8 fold higher risk of death compared to AKI caused by other etiologies ^6^.

Current understanding of the pathogenesis of S-AKI mainly includes alterations in renal microvascular circulation, adaptive immune responses to exacerbated inflammation, and reprogramming of metabolic pathways ^6,7^. First, altered microvascular blood flow impairs tissue oxygenation, which in turn leads to multiple organs’ dysfunction. In the kidney tissue, glomerular shunting and constriction of the efferent arteriole result in decreased intraglomerular pressure and subsequently reduced glomerular filtration rate and urine output ^4,6,8^. Increased vascular permeability and leakage can also cause interstitial edema and increase oxygen diffusion distant to the tubular cells. Endothelial activation promotes rolling and adhesion of leukocytes and platelets, resulting in an increased risk of thrombi formation and flow continuity alterations. Second, tubular endothelial cells are equipped with pattern recognition receptors (such as Tolllike receptors, TLRs), which can receive signals from sepsis-induced inflammation, pathogen-associated molecular patterns (PAMPs), or damage-associated molecular patterns (DAMPs). These result in a cascade of downstream signals and increase the synthesis of proinflammatory cytokines, reactive oxygen species, and oxidative stress that damage renal tubular cells ^4,8,9^. Lastly, the metabolism and energy state of renal tissue is reprogrammed during sepsis to conserve and reprioritize energy spending for the tubular cells to recover from damage ^8,10,11^. However, the global changes of the diseased kidneys after S-AKI at the molecular level remain largely unknown.

The human gastrointestinal (GI) tract harbors over 100 trillion microbial cells from nearly 2,000 bacterial species ^12,13^. These GI tract-associated microbial organisms are often referred to as the gut microbiome ^14^. Traumatic gut injuries and/or intestinal inflammation leading to the breakdown of the epithelial barrier of the GI tract can cause a so-called ‘leaky gut,’ allowing translocation of bacteria and toxins from the intestinal lumen to the mesenteric lymph and the circulation system, thus driving systemic inflammation ^14–16^. Indeed, previous studies have shown the detection of gut bacterial DNA fragments in the blood of CKD patients ^17,18^. From the bloodstream, gut microbes may gain access to local tissues or distal organs, amplifying existing injury and leading to persistent injuries. Infection of bladder urothelium has been reported with uropathogenic *Escherichia coli*, a gut-originated bacteria that can replicate intracellularly, causing recurrent urinary tract infections and cystitis ^19,20^. Additionally, striking data from recent studies indicated that the respiratory tract extracellular pathogen *Streptococcus pneumoniae* could translocate to heart tissues and replicate intracellularly, where it transforms into immunoquiescent biofilm, induces cardiac microlesions, and exacerbates community-acquired pneumonia ^21,22^. However, a prevailing paradigm in renal physiology is the assumption that the kidney is sterile ^23^. Whether gut microbes can invade and colonize kidney is still an unanswered question. Therefore, we constructed an S-AKI mouse model using the cecal ligation and puncture (CLP) procedure. CLP is considered to be a gold standard clinical model ^24^ as it allows the release of fecal components into the peritoneal cavity and circulatory system that induces exacerbated inflammatory responses. This procedure mimics gut injury-triggered severe sepsis in humans. Hence, it is considered to be more clinically relevant when compared to the endotoxic model ^25,26^. In the current study, we analyzed the global proteome and phosphoproteome level changes of kidney tissues at day 2 and day 7 after CLP. With the results we obtained, we can better characterize the molecular mechanisms underlying the development of S-AKI and its transition to CKD.

## Experimental Procedures

### Ethics statement, mouse models, and CLP surgery

Male C57BL/6J mice weighing about 6-8 weeks and 20–25 g were obtained from the Jackson Laboratories (Bar Harbor, ME). Cecal ligation and puncture procedure was performed by using an established protocol described previously ^27^. Briefly, the abdomen of the mice was shaved and disinfected after anesthetizing the mice. Under aseptic conditions, a 1-2 cm midline laparotomy was performed to expose the cecum with the adjoining intestine. The cecum was tightly ligated with a 6.0 silk suture at its base below the ileo-cecal valve (1 cm distance from the distal end of the cecum to ligation point) and was perforated twice with a 19 gauge needle on the same side of the cecum. The cecum was then gently squeezed to extrude a small amount of feces from the perforation sites. The cecum was returned to the peritoneal cavity and peritoneum and skin were closed. Mice were sacrificed at 0, 2, or 7 days after the procedure. Kidney tissues were collected for various analyses. All the above animal experiments were approved by the Institutional Animal Care and Use Committee at the University of Pittsburgh.

### Determination of Serum Creatinine

Serum was collected from mice at 1 day, and the serum creatinine level was determined using the QuantiChrom creatinine assay kit according to the protocols specified by the manufacturer (BioAssay Systems, Hayward, CA). The level of serum creatinine was expressed as milligrams per 100 ml (dl) as described previously ^28^.

### Histology staining, western blot and ELISA analysis

Paraffin-embedded mouse kidney sections (~3 μm thickness) were prepared and then stained with Periodic acid–Schiff (PAS) reagents following standard protocols ^29^. For immunoassays, kidney tissues were lysed with radioimmunoprecipitation assay (RIPA) buffer containing 1% NP-40, 0.1% SDS, 100 μg/ml PMSF, 1% protease inhibitor cocktail, and 1% phosphatase I and II inhibitor cocktail (Sigma) in PBS on ice. The supernatants were collected after centrifugation at 13,000×g at 4°C for 15 minutes. Protein expression was analyzed by Western blot analysis as described previously ^30^. The primary antibodies used were as follows: anti-FADD (05-486; Millipore, Billerica, MA); anti-NGAL (ab63929), anti-TNF-α (ab1793; Abcam, Cambridge, MA); anti-Bad (#9268; Cell Signaling Technology, Danvers, MA); anti-Gasdermin D (sc-81868; Santa Cruz Biotechnology, Santa Cruz, CA); and anti-α-tubulin (T9026; Sigma, St. Louis, MO). For ELISA analysis, kidney samples were processed for TNF-α expression using the Mouse TNFα Quantikine ELISA Kit (R&D Systems MTA00B). Samples were diluted 1:5 in reagent diluent prior to plating. Assay was performed according to manufacturer’s instructions.

### Global proteomic and phosphoproteomic sample preparation

Mouse kidneys were homogenized in PBS with HALT protease inhibitor cocktail (Pierce) in a bead beater. After collecting soluble homogenate, the tissue debris was re-suspended in 2 x SED lysis buffer (4% SDS, 50 mM EDTA, 20 mM DTT, 2% Tween 20, 100 mM Tris-HCl, pH 8.0) and sonicated to further extract membrane-bound proteins. The homogenate from both PBS and SED treatment were combined (~500 μg proteins) and digested using the homemade STrap method. ~50 μg digested peptides were directly desalted using spinnable StageTip protocol ^31^. To analyze the phosphoproteome of the kidney tissue, ~450 μg digested peptides were processed with phosphorylation enrichment using TiO2 beads as described previously ^32^. Peptides were desalted using StageTip as described, lyophilized and stored in −80°C until futher analysis ^33^.

### LC-MS/MS analysis

The LC-MS/MS analysis was carried out using an Ultimate 3000 nanoLC coupled to Q Exactive mass spectrometer (Thermo Scientific). Peptides were first loaded onto a trap column (PepMap C18, 2 cm x 100 μm x I.D.; Thermo Scientific) and then separated by an in-house packed analytical column (C18 ReproSil, 3.0 μm, Dr. Maisch GmbH; 19 cm x 75 μm I.D.) using a binary buffer system (buffer A: 0.1% formic acid in water; buffer B: 0.1% formic acid in acetonitrile) with a 220-min gradient (2-35% buffer B over 180min; 35-80% buffer B over 10min; back to 2% B in 5 min for equilibration after staying on 80% B for 5 min). MS data were acquired in a data-dependent top-10 method with a maximum injection time of 20 ms, a scan range of 350– 1700 Da, and an AGC target of 1e6. MS/MS was performed via higher energy collisional dissociation fragmentation with a target value of 5e5 and a maximum injection time of 100 ms. Full MS and MS/MS scans were acquired at a resolution of 70,000 and 17,500, respectively. Dynamic exclusion was set to 20s.

### Protein identification and quantitation

Protein identification and quantitation were performed using the MaxQuant-Andromeda software suite (version 1.6.5.0) with most of the default parameters ^34^. A mouse database (17,038 sequences; Reviewed only; version July 2019) was used for the database search. For global proteome analysis, the following parameters were applied: 10 ppm and 20 ppm mass tolerances for precursor and fragments, respectively; trypsin as enzyme with two missed cleavage sites; protein N-terminal acetylation and methionine oxidation as variable modifications; cysteine carbamidomethylation as a fixed modification; peptide length with at least 7 amino acids. For phosphoproteome analysis, phosphorylation at serine, threonine, and tyrosine was set as an additional variable modification. False discovery rate (FDR) was set at 1% for both proteins and peptides. The cutoff of phosphosite probability estimated by MaxQuant was required to be 0.75 or higher. Further bioinformatics analyses such as Hierarchical clustering, Principal Component Analysis (PCA), t-tests, correlation, and volcano plots were performed in the Perseus environment (version 1.6.1.3). The MaxQuant output result (proteinGroups.txt) was first filtered to exclude those ‘Only identified by site, ‘Potential contaminant’, and ‘Reversed’ hits, and then log2-transformed.

Gene Ontology (GO) and Kyoto Encyclopedia of Genes and Genomes (KEGG) pathway analyses were performed using DAVID Bioinformatics Resources 6.8 (https://david.ncifcrf.gov/home.jsp). For protein network analysis, the StringApp was employed in the Cytoscape environment (version 3.8.0) ^35^. The interaction score was set to 0.9, the highest confidence cutoff, to retrieve potential interactions. The ‘Load Enrichment Data’ option was enabled during this process to retrieve functional enrichment (minimum significance threshold FDR 0.05) for the STRING network.

### Experimental Design and Statistical Rationale

The mouse CLP experiments and biospecimens at each time point were collected in at least six biological replicates. Western blot and biochemical assays were performed for all the collected samples. Proteomic and phosphoproteomic experiments were performed for six samples of each time point. All tissues were lysed and digested in parallel. For global quantitation of all proteomics and phosphoproteomics data, the files were processed in MaxQuant software in the same batch as well. Multi-sample variation test was determined by ANOVA in the Perseus environment corrected by the Permutation FDR 0.05. Pair-wise Student’s t-test was performed following a similar procedure, and the p values were corrected as well with Permutation FDR 0.05. Unsupervised clustering analyses used Euclidean as distance and average as linkage for both column and row clustering.

## Results

### Characterization of kidney injury after S-AKI

Given that S-AKI is the most common complication associated with sepsis in the clinic, we aimed to explore its pathogenesis in detail. We created a moderate S-AKI mouse model in which the mice would survive to progress to chronic kidney disease. CLP surgery was performed by placing a 2-0 silk ligature at 1 cm from the cecal tip as described previously (**Figure 1A)**, with mice sacrificed at day 2 or day 7 ^24^. Only 1 out of 20 mice that went through the procedure died before the entire experiments were finished, indicating we did not induce severe sepsis. Next, we examined if the CLP procedure successfully induced AKI using rapid elevation of serum creatinine (SCr) as an established biomarker of AKI ^36^. We observed that SCr was significantly increased to an average of 0.33 mg/dl at 2 days after the CLP procedure, a nearly 3-fold increase compared to the sham control group. By 7 days after CLP, SCr returned to a relatively low level (**Figure 1B**). We also investigated the expression of neutrophil gelatinase-associated lipocalin protein (NGAL; also known as lipocalin 2, Lcn2), a validated predictor for kidney damage. Under ischemic, septic, or post-transplant AKI, NGAL’s expression in kidney tubules is often rapidly increased ^37^. NGAL expression increased dramatically in renal tissues at day 2 after CLP compared to the sham control, with a reduction by day 7 (**Figure 1C** and **1D**), indicating that CLP caused a definite injury to the kidneys. Consistent with this, periodic acid-Schiff (PAS) staining illustrated cell vacuolation, necrosis, and cellular debris present near blood vessels in the diseased kidneys at day 2 after CLP (**Figure 1E**), which are consistent with previous observations of septic kidneys ^38^. Little to no differences in histological changes were observed between day 2 and day 7 after CLP. In contrast to S-AKI, ischemia-reperfusion associated AKI usually causes more profound histological changes, as reported by our previous study ^39^.

**Figure 1.**
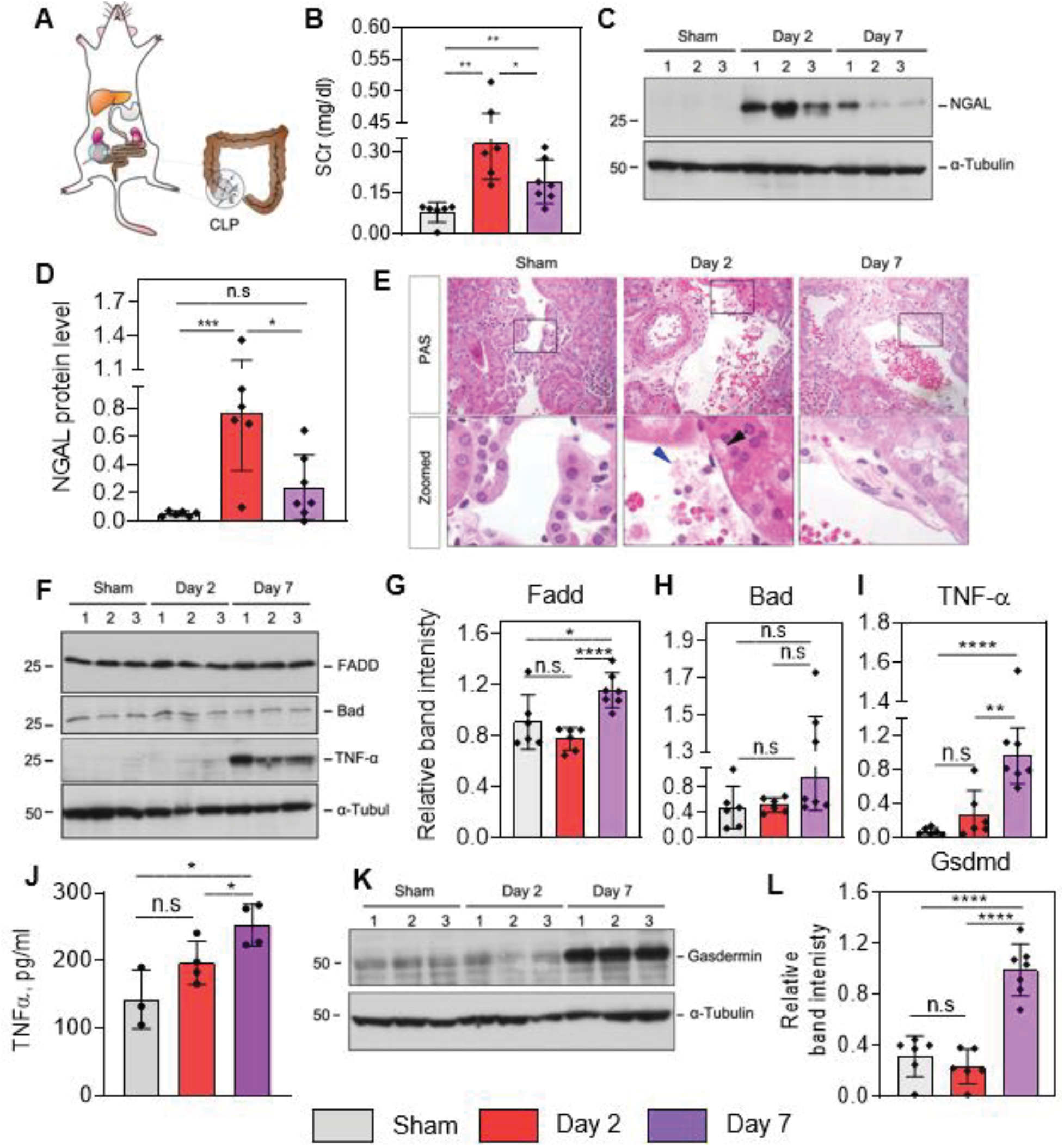
Cecal ligation leads to sepsis, kidney inflammation and pyroptosis in a time dependent manner. **(A)** Schematic diagram illustrating the construction of the moderate CLP mouse model. **(B)** Measurement of serum creatinine (SCr) levels from control mice (sham), day 2, and day 7 post-CLP surgery (n = 6-7). **(C)** Western blot analyses of renal expression of NGAL protein in sham control and injured kidneys after CLP at day 2 and 7. Numbers (1-3) indicate each animal in a given group. (**D**) Quantitative plots of (**C**) with more data points included. (**E**) Representative micrographs showing kidney morphology at day-2 and day-7 after CLP. In the enlarged boxed areas, the black arrow indicates typical tubular cell vacuolation after CLP, and the blue arrowhead indicates cellular debris and possible migrated microbial. Scale bar, 50 μm. (**F**) Western blot analyses of FADD, Bad, and TNF-α protein abundance in the kidneys at day 2 and 7 after CLP compared with the sham control. Numbers (1-3) indicate each animal in a given group. (**G-I**) Quantitative data of (**F**) with more data points. (**J**) ELISA results of TNF-α level in the kidneys at day-2 and day-7 after CLP. (**K**) Western blot analyses of Gasdermin D abundance in the kidneys at day 2 and 7 after CLP compared with the sham control. (**L**) Quantitative data of (**K**) with more data points.

It has been known that cell apoptosis and inflammatory cell infiltration are two major cellular processes in the development and progression of AKI ^40^. To investigate this, we examined two cell apoptosis-related proteins, Bcl2-associated agonist of cell death protein (Bad) and FAS-associated death domain protein (Fadd) ^40,41^. Bad appeared to remain unchanged (p > 0.05) across the three time points, whereas Fadd showed mild increase at day 7 in comparison to sham control (**Figure 1F** through **1H**). Consistently, the expression of tumor necrosis factor-α (TNF-α) displayed a similar trend to Fadd in the S-AKI kidneys by western blot (**Figure 1I**) and enzyme-linked immunosorbent assay (ELISA) (**Figure 1J**). As necrosis was observed in our histological analysis, we tested for the activity of the effector protein of pyroptosis, Gasdermin D (Gsdmd) ^42^. This pathway has only recently been associated with kidney injury ^43^. Gasdmd was significantly increased in the injured kidneys at day 7 after CLP (**Figure 1K, 1L**). Collectively, these data strongly suggest that a delayed biological response to septic injury in kidney dominates the transition from AKI to CKD.

### The global proteomic view of the septic kidneys

To gain an unbiased understanding of the underlying molecular determinants that modulate S-AKI, we analyzed the global profile of the renal tissue proteomes of mice at day 0, 2 and 7 post-CLP following a label-free quantitative approach (**Figure 2A**). The global kidney proteome was analyzed using a 220-min single-run LC-MS/MS approach to balance the throughput and proteomic depth. Collectively, 2,119 proteins were quantified with a less than 1% false discovery rate (FDR) across all three groups (**Supplementary Table S1**). The correlation between biological replicates (i.e. different mice) within the same group was high (R^2^ = 0.95 ± 0.02, n = 45) (**Figure 2B**), which validated the reliability of our CLP model and the reproducibility of the sample processing approach. We also noticed that the correlation between different groups was relatively high as well (R^2^ = 0.93 ± 0.02, n = 108). These data indicate that the CLP procedure induced low-to mid-grade sepsis, likely only allowing a minimal amount of feces to extrude from the perforation site. Interestingly, the overall unfiltered proteomics data enabled the unsupervised classification of the kidneys according to their global profiles, which clustered day-2 mice tightly together while control and day-7 samples were intermixed (**Supplemental Figure S1**). The data suggested that the kidney global proteome has undergone systematic changes after moderate septic injury, and may have reshaped back closer to healthy kidneys a week after CLP.

**Figure 2.**
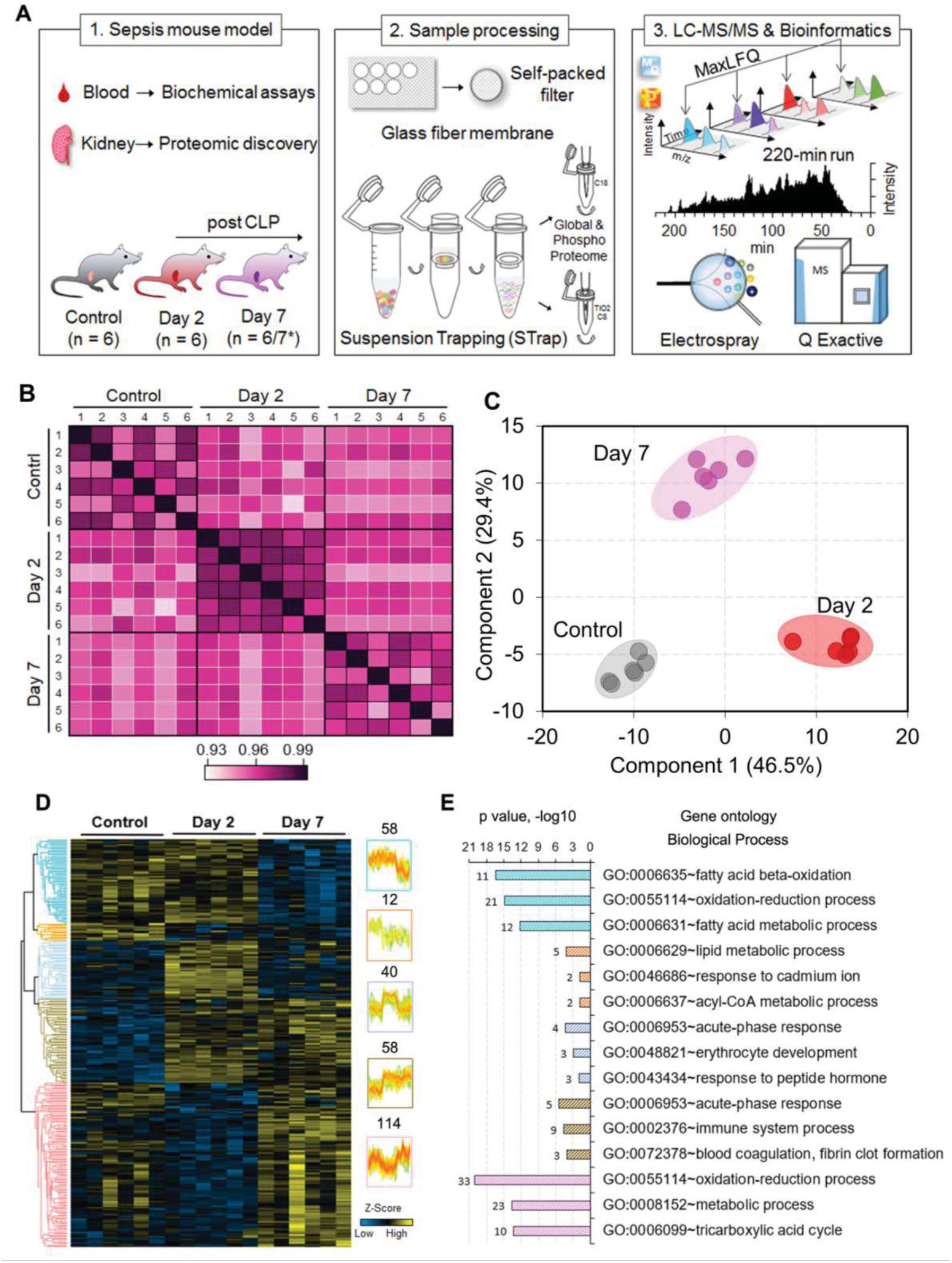
Differential proteomic profiles define acute versus chronic kidney injury. (**A**) Experimental workflow for analyses of proteome and phosphoproteome of mouse kidneys. (**B**) Correlation of kidney proteome profiles between different groups of mice. Color scale represents R^2^ values. (**C**) and (**D**) PCA lot and heatmap of ANOVA significant proteins (Permutation FDR 0.05) among the three groups. LFQ intensity of represented proteins was z-scored and plotted according to the color bar. Five clusters of proteins with different patterns of abundance profile are highlighted in the heatmap. The numbers indicate the number of proteins in each cluster. (**E**) Top three enriched GO biological process terms in each cluster of proteins are plotted with their names and significance (*p* value). The number of matched proteins in each category is listed next to the bar graph.

To define the proteins that were differentially regulated in response to septic injury, and determine underlying modulatory pathways, we performed a multiple-variations test (ANOVA). We first filtered the overall proteome to require that proteins be quantified in at least 4 out of 6 replicates in one of the groups, which led to 1,520 high quality proteins. The ANOVA analysis resulted in 282 significant proteins among the three groups (Permutation FDR 0.05) (**Figure 2D, Supplementary Table S2**), which was further verified by their distinct segregation in a PCA plot (**Figure 2C**). Gene Ontology (GO) enrichment analysis indicated that these proteins were generally involved in the oxidation-reduction process, metabolic process, and fatty acid beta-oxidation. While acute-phase response proteins and proteins that are associated with the immune system process were largely increased upon septic injury, oxidation-reduction, as well as metabolic process proteins, showed varied profiles (**Figure 2E**). This data highlighted the complex and dynamic host response to the septic infection. By examining the subcellular localization of the ANOVA significant proteins, we found that nearly 44% of them (124 out of 282) were enriched in “mitochondrion” (*p* = 8.14 x 10^−57^) (**Supplemental Figure S2**). Of note, around 21% of mitochondrial-associated proteins were identified in a global kidney proteome ^44^, and less than 10% in the overall mouse proteome (17,034 Reviewed UniProt proteins). Among the mitochondrial proteins, 46 (37%) of them were annotated as mitochondrial inner membrane proteins, such as ATP synthase (e.g., Atp5j, Atp5l, and Atp5o), NADH dehydrogenase (e.g., Ndufb1/3/4/7/9/12/13) and cytochrome c oxidase (e.g., Cox2 and Cox4i1). This finding is consistent with the biological process enrichment result, and suggests that mitochondrial dysfunction is strongly associated with the decline of kidney integrity and hemostasis ^45^, and in our case, during sepsis ^46^.

### Septic injury induces dynamic changes in kidney phosphoproteome

To gain more insights into the temporal regulation and functional changes of kidney proteins during S-AKI, we performed phosphoproteome analysis to the same set of tissue samples used for the global proteome analysis (as shown in **Figure 2A**). In total, 4,745 phosphosites were identified from our study, of which 2,950 were unambiguously localized with high confidence (localization score > 0.75), corresponding to 1,279 non-redundant phosphoproteins. These phosphorylation events were used for downstream analysis and are accessible online (**Supplementary Table S3**). The majority (64%) of the 1,279 proteins were exclusively identified from the phosphoproteomics experiments (**Supplemental Figure S3A**), implying the specificity of the TiO2-based phosphorylation enrichment process. In the context of the reproducibility of phosphoproteome measurements, a much lower correlation was seen both between biological replicates (average R^2^ = 0.82) and different groups (average R^2^ = 0.79) (**Supplemental Figure S3B**) as compared to those of global proteomes. These data suggest that different phosphorylation patterns exist, as indicated by a previous tissue phosphoproteome study ^47^, and temporal phosphoproteomic changes in response to microbial infection are more dynamic than global proteomic changes.

To base our analyses on high quality data, we filtered the phosphorylation data set to exclude samples that contained less than 1,000 quantifiable phosphosites, and phosphosites that were quantified in less than 3 replicates, which resulted in 1,856 phosphosites. Subsequent ANOVA analysis resulted in 388 significant phosphosites among the three groups (p < 0.05) (**Supplementary Table S4**), and further p-value correction (Permutation FDR 0.05) resulted in 22 phosphosites, as presented in **Figure 3A**. Overall, the majority of the kidney phosphorylation events drastically decreased at day 2 post CLP and gradually returned to basal level at day 7 (**Supplementary Figure S4 and Figure 3, F-K**), indicating a declined energy state of the renal tissue upon acute infection and dynamic host response to septic injury. Enrichment analysis indicated that phosphoproteins associated with RNA splicing and mRNA processing were significantly over-represented (p < 10^−13^) (**Figure 3B**). These include RNA binding motif protein families (e.g., Rbm10/17/25/39/xl1), serine/arginine-rich splicing factors (e.g., Srsf1/9) and serine/arginine repetitive matrix proteins (e.g. Srrm1/2). Alternative RNA splicing regulates host immune response in a variety of viral and bacterial infection conditions ^48^, and RBM proteins have been found to modulate apoptosis during infection in addition to splicing ^49^. As an example, the phosphorylation of Srrm2 has been shown to modulate human immunodeficiency virus (HIV-1) replication and release ^50^. Srrm2 is highly phosphorylated, with over 250 known phosphorylation sites listed in the UniProt Knowledgebase. In our study, we identified 79 of them, similar to a previous study ^51^. About 26 sites were differentially phosphorylated upon sepsis injury, including a highly significant one at serine 1229/1230 (ANOVA p < 0.0001). Its modulatory role in polymicrobial infection deserves further investigation.

**Figure 3.**
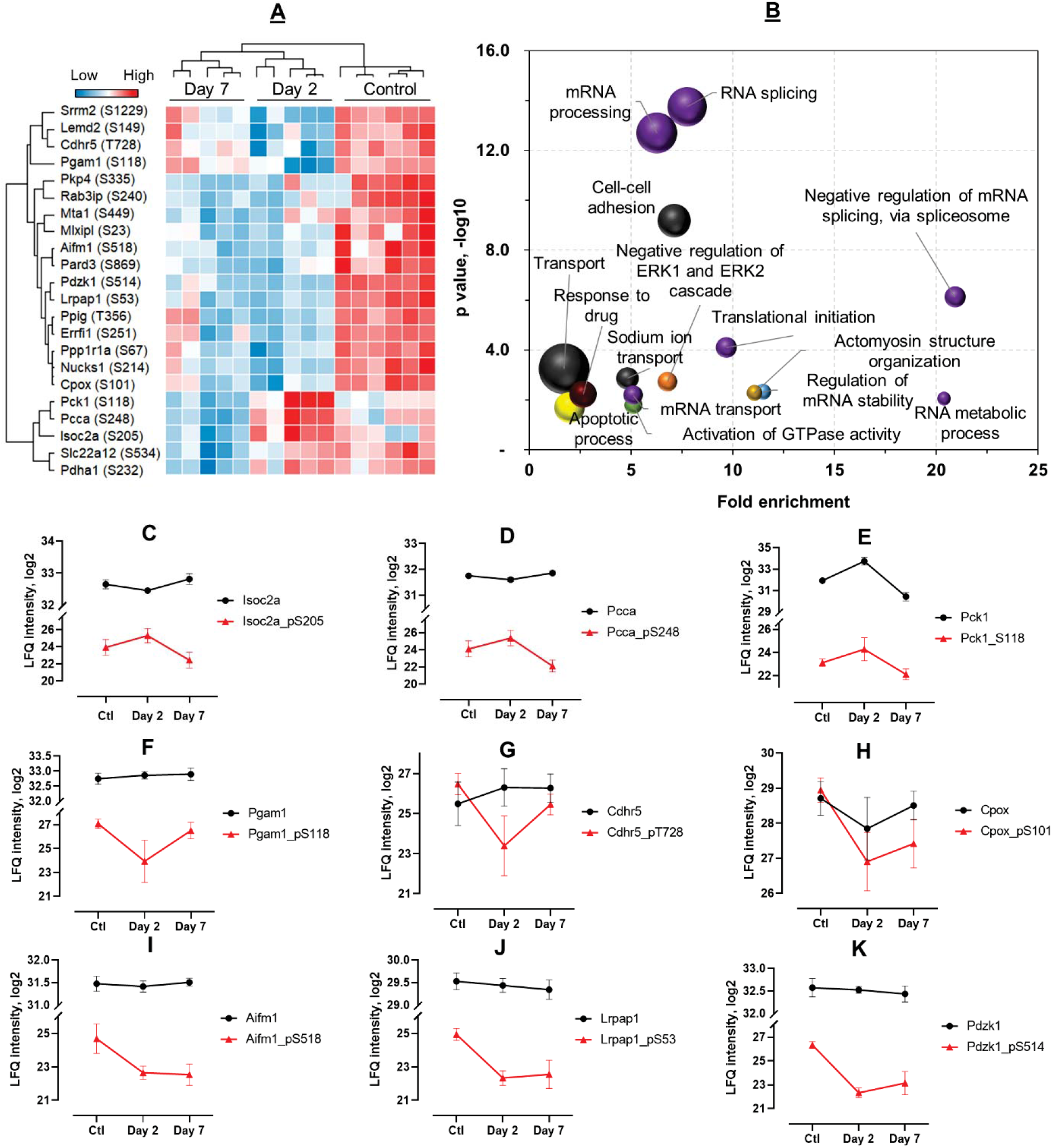
Quantitative phosphoproteome analysis of kidneys after septic injury. (**A**) Heatmap of representative phosphosites significantly altered among the three groups (ANOVA, Permutation FDR 0.05). (**B**) Enrichment analysis of biological process of ANOVA significant phosphoproteins. The top-15 terms are plotted. Bubble size depicts the number of protein in the category, ranging from 3 to 42. (**C-K**) Intensity plots of representative phosphosites (red line) and their unmodified forms (black line). Ctl, control.

While most of the phosphorylation events were reduced after acute septic injury, a small number of them remained high at day 2 (**Figure 3, C-E**). These include isochorismatase domaincontaining protein 2A (Isoc2a, S205), mitochondrial propionyl-CoA carboxylase alpha chain (Pcca, S248), and phosphoenolpyruvate carboxykinase (Pck1, S118). The identified phosphorylation at S205 in Isoc2a is novel, therefore its role in S-AKI needs further investigation. To distinguish whether the increased phosphorylation levels of these proteins at day 2 were due to changes in their total protein abundance, we compared their phosphorylation intensities with their respective total protein levels (**Figure 3, C-E**). Interestingly, the expression levels of both Isoc2a and Pcca remained stable among the three groups (p > 0.05), whereas their phosphorylated forms changed significantly (up-regulated at day 2 and down-regulated at day 7; p < 0.05), indicating a true upregulation of their phosphorylation at day 2. As for Pck1, similar abundance changes were seen in both its phosphorylated and unmodified forms, suggesting its regulation in response to septic injury was partly due to changes in its total protein level. Pcca is a mitochondrial enzyme involved in amino acid catabolism ^52^ and mitochondrial oxidative phosphorylation. In *C. elegans*, its mutation significantly reduced mitochondrial energy metabolism and increased oxidative stress ^53^, and previous studies have demonstrated that it is strongly linked to the genetic disease propionic acidemia ^53^. Phosphorylation of Pcca at site S248 has been identified by proteomic screening from various mouse tissues previously ^54,55^, however, the phosphorylation/dephosphorylation of this site in response to polymicrobial infections and its association with mitochondrial metabolism in kidneys is unknown.

An interesting class of proteins that was identified in our analyses is the solute carrier (SLC) proteins, a superfamily of membrane-bound transporters with nearly 400 members ^56^. They regulate ion transport, waste removal, and many other essential physiological functions, and have increasingly been targeted for therapeutic invention ^57^. However, their involvement in septic injury has not been well studied. In our global and phosphoproteomic studies, 72 SLC proteins as well as 113 phosphosites were identified (**Supplemental Table S2 and S4**).). The majority of SLC proteins identified in our study did not show significant differences upon septic injury, and even if changed, the fold difference were minimal (**Figure 4B**). The significantly (AVANO, p < 0.05) changed SLC proteins and/or their phosphorylated forms are shown in **Figure 4A**. The Band 3 anion transport protein (Slc4a1) increased dramatically (FC > 4) during acute septic infection at day 2, whereas its phosphorylated form (pS18) did not change (**Figure 4C**). On the other hand, the unmodified form of the solute carrier family 22 member 12 (Slc22a12) did not show significant alterations, but its phosphorylation at S534 was maintained at a similar level at day 2 and then down-regulated at day 7 (**Figure 4D**). Phosphorylation of Slc22a12 (also known as Urat1) has been shown to assist in the reabsorption of uric acid via other organic anion transporters ^58,59^. Lastly, both the unmodified and phosphorylated forms of the solute carrier family 22 member 2 (Slc22a2) were changed in a similar trend after septic injury (**Figure 4E**). These data indicate that the homeostasis of ions and small molecules in the kidney are broadly altered after sepsis induction.

**Figure 4.**
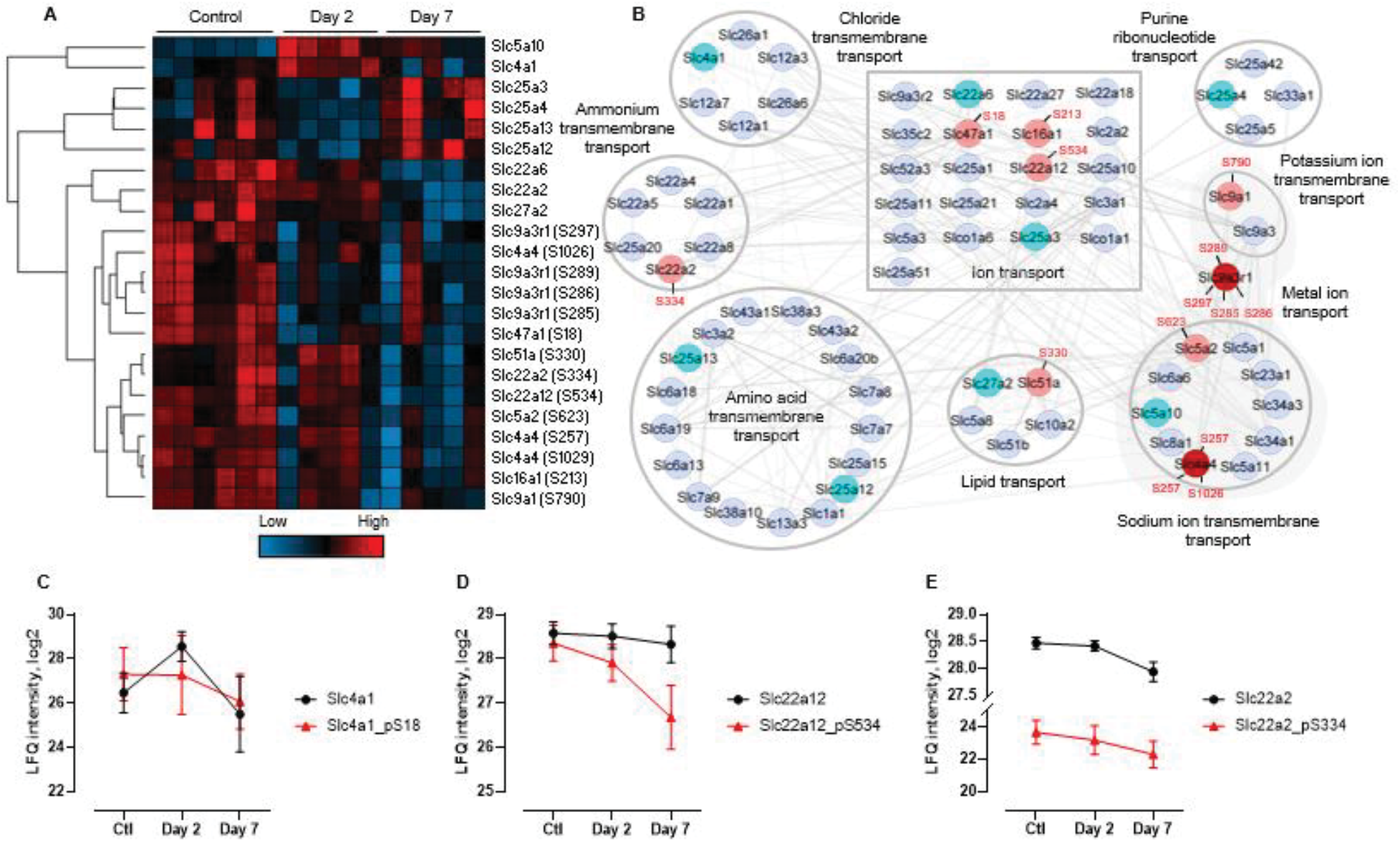
Quantitative analysis of SLC proteins. **(A)** Heatmap of ANOVA significant (p < 0.05) proteins and phosphosites. (**B**) Protein network and functional enrichment clusters. The network was built based on all the 72 identified SLC proteins as input using the String App in Cytoscape software. nodes colored in light blue are non-differential (FC < 1.5) proteins; in cyan are significantly different proteins; in pink and red are differential phosphoproteins. The protein clusters were annotated based on the Enrichment Data, an embedded function in Cytoscape. (**C-E**) Profiles of representative SLC proteins across the three groups with phosphosite intensities in red line and unmodified forms in black.

### Discovery-based studies recapitulate signature proteins for S-AKI prognosis and disease progression

Next, we sought to examine signature proteins that correlate with the severity of S-AKI. We performed pairwise comparisons between control and septic kidneys at day 2 or day 7 to highlight significant marker proteins (**Figure 5, A-B; Supplementary Table S5**). In our data, the known AKI marker protein NGAL dramatically increased (> 64-fold; p < 0.001) at day 2 and gradually lowered down to near basal level (p > 0.05; **Figure 5C**). This finding was consistent with the immunoblot analysis (**Figure 1, C-D**), highlighting the value of our MS-based approach for kidney disease biomarker discovery. In addition to NGAL, we also identified other marker proteins and proteins that have not been previously associated with kidney injury. Hydroxymethylglutaryl-CoA synthase 2 (Hmgcs2), a key enzyme of ketogenesis mediating energy generation from lipids when carbohydrates are deprived, was increased in day-2 kidneys by more than 80-fold (**Figure 5D**). Hmgcs2 has been indicated to be up-regulated in both Type II diabetic kidneys and Type I diabetic heart ^60,61^, suggesting enhanced ketone body production and activated ketogenesis in these diseased organs. Serine protease inhibitor A3N (Serpina3n) was also significantly increased at day-2 by nearly 60-fold (**Figure 5E**). Serine protease inhibitors are usually found to be increased during acute inflammation and act by modulating protease activities ^62,63^. Recently, Serpina3n was found to be elevated in the urine of rats with early AKI-to-CKD transition and relocated from cytoplasm to apical tubular cell membranes ^64^. Other proteins such as serum amyloid A-1 and A-2 protein (Saa1 and Saa2) and serum amyloid P-component (ApcS), proteins that were shown to be increased in diabetic kidneys and chronic renal failure ^65,66^, were also markedly increased in day-2 post-CLP kidneys (**Figure 5, F-H**).

**Figure 5.**
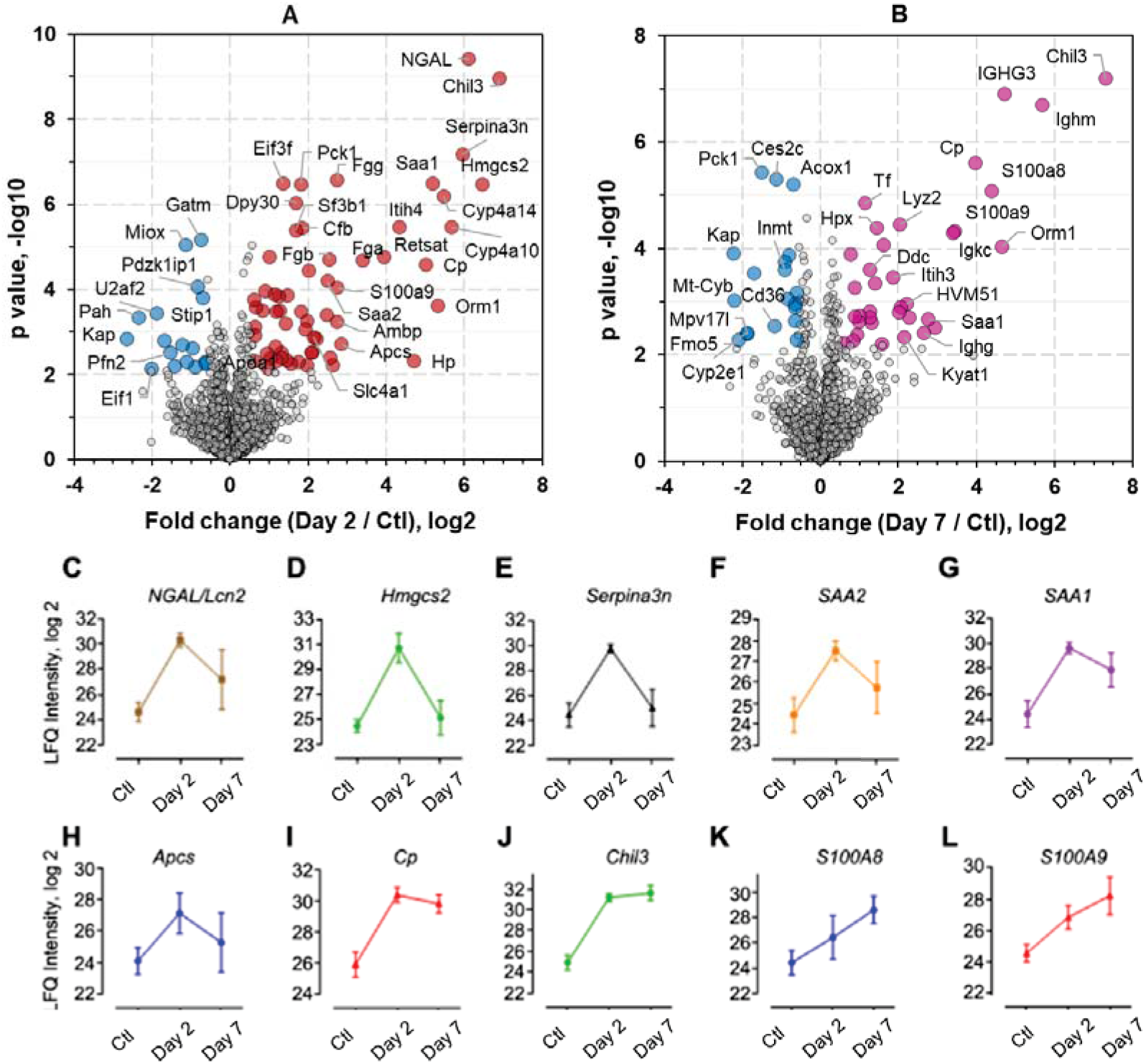
Kinetic changes of sepsis-induced kidney injury markers. **(A-B)** Volcano plot of pairwise comparison between the kidney proteomes of day 2 or day 7 post-CLP versus control. Student t-test (Permutation FDR 0.05) was used for the analyses. Up- and down-regulated proteins by 1.5-fold were colored in red/purple and blue, respectively. Gene name of represented proteins were indicated in the plots. **(C-L)** Abundance plots of representative proteins in response to septic injury. Mean values and error bars from six replicates of each group were shown. Ctl, control.

Of note, a number of proteins showed consistent up-regulation at both day 2 and day 7 post-CLP. Ceruloplasmin (Cp) is a copper-containing ferroxidase that oxidizes ferrous iron (Fe^2+^) to its nontoxic ferric iron (Fe^3+^) form. Its expression was shown to be increased in aging mice or mice consuming high-calorie diet ^67^, suggesting the increased amount of oxidative damage in these mice required enhancement for its antioxidant activity. The urinary Cp level was suggested to be a biomarker for CKD in sickle cell disease patients ^68^. Cp levels in our day-2 and day-7 mice post-CLP were elevated by 32- and 15-fold, respectively (**Figure 5I**), suggesting increased oxidative stress in kidneys post-sepsis. Meanwhile, S100A8 and S100A9, DAMPs molecules that mediate pro-inflammatory responses *via* TLR4 binding ^69^, were previously indicated to be increased in plasma and associated with septic shock ^70^. In our proteome analysis, the increase was significant at both time points for S100a9, whereas it was significant only at day 7 for S100a8 (**Figure 5, K-L**), suggesting the formation of the active dimerized calprotectin was gradually increased in kidneys after septic injury ^69^.

Among the proteins that were significantly increased post-CLP, we found the upregulation of chitinase-like protein 3 (Chil3) to be most striking, by almost 120-fold at day 2 and more than 150-fold at day 7 (**Figure 5J**). Chil3 is a member of the mammalian chitinase family that lacks the chitinase activity though it is highly secreted by macrophages and neutrophils ^71,72^. Similar to the function of S100 proteins, chitinase-like proteins have also been shown to be mediators of inflammation and antimicrobial responses in multiple models of disease such as asthma and pulmonary fibrosis as well as involved in tissue remodeling and wound healing ^73–75^. In previous work in kidney disease models, a urinary proteomic study has shown that chitinases-like proteins could be candidate biomarkers for S-AKI ^75^. Recently, it has been demonstrated to be largely increased in urine from clinical AKI patients ^76^. Another study also showed renal Chil3 to be up-regulated at 6 hours and 24 hours after intraperitoneal injection of LPS into mice ^77^. Taken together, our study recapitulated previously described sepsis-related modulator proteins and provided a valuable resource of candidate targets to future studies of sepsis diagnosis and therapeutics.

### Molecular-level understanding of AKI-CKD transition

As described above, our data have shown distinct pathology of the septic kidneys at the acute and chronic stages by analyzing them at day 2 and day 7 post-septic injury, respectively. Therefore, a direct comparison between these two phases would lead to insights into how the disease progresses after acute septic injury. Following a similar t-test (FC ≥ 1.5; Permutation FDR 0.05), we identified 38 and 56 up- and down-regulated proteins, respectively, from day-7 mice in comparison to day-2 mice (**Supplementary Table S6**). Interestingly, although the functional enrichment analysis did not suggest particular biological process terms enriched among the up-regulated proteins, likely due to the low number of input proteins, we noticed several immunoglobulin proteins (such as Ighm, Ighg, Igkc, IGHG3, and HVM51) were significantly increased. This suggested that while recovering from the acute phase of septic-induced kidney injury, there may be an initiation of an adaptive immune response in renal tissue progressing toward chronic kidney disease. Indeed, a switch from innate to adaptive immunity has been reported to mediate the transition from acute septic kidney injury to prolonged chronic disease ^78,79^, and some of these immunoglobulins can be used to develop biomarkers for CKD in the future ^80^.

To obtain a broader view of the dysregulated pathwaysduring S-AKI-CKD transition, we slightly lowered the fold-change threshold (≥ 1.2) to capture more differential proteins (202 in total; **Figure 6A**) and build a signaling network (see details in Experimental Procedures). The analysis with the highest confidence score (cutoff = 0.9) resulted in 396 interactions and 126 nodes (**Figure 6B**), the majority (56%) of which were associated with small-molecule metabolic processes (FDR = 3.35e-48), such as lipids, nucleotides, alcohol, and other fatty acids. Abnormal lipid metabolism has been associated with diabetic nephropathy and other kidney dysfunctions ^81,82^; lipid deposition contributes to atherosclerosis and cardiovascular diseases commonly seen in CKD patients ^83^. The network analysis also indicated several interesting protein clusters such as oxidoreductase complex, peroxisome, and cytochrome P450 (CYP) family proteins (**Figure 6B**), which may play important roles in the AKI-CKD transition. Peroxisomes and oxidoreductases invovles in regulating oxidative stress caused by bacterial and/or viral infections ^84^. In our data, 18 of the 21 proteins annotated as peroxisome proteins showed down-regulation in day-7 kidneys. In contrast, all the 12 oxidoreductase complex associated proteins were up-regulated in day-7 kidneys. The role of oxidoreductases in the progression of the disease stages of septic kidneys remains to be identified. CYP proteins are known to metabolize a wide array of small molecules such as fatty acids, alkanes, and xenobiotic drugs. Recent studies have shown their involvement in microbial infections ^85,86^. In our analysis, five CYP members (2A4, 2A5, 2E1, 4A10, and 4A14) were found to be significantly down-regulated in day-7 kidneys compared to day-2. For example, the expression of Cyp4a10 was dramatically increased after acute infection (at day 2) and declined back to the basal level at chronic stage (at day 7) (**Figure 6C**). It has been noted that CYP enzymes can be selectively regulated in different states of inflammation by a multitude of cytokines released after infection or inflammation ^85^. In hepatic and extrahepatic tissues, chronic inflammation is usually associated with downregulation of CYP enzymes ^85^. Hence, our quantitative data of CYP proteins were in line with previous studies and suggest that they could also be used as signatures for kidney disease severity and/or classification. Notably, carboxylesterase 1C (Ces1c), which aids in detoxification of xenobiotics and is responsible for poor pharmacokinetics of the anti-virus drug remdesivir ^87^, was also mapped in our cluster analysis. Ces1c has been shown to be significantly decreased in macrophages after LPS treatment ^88^, which was consistent with its changes in our data of septic kidneys (**Figure 6D**). Together, our proteomic data provides an unbiased view of how kidney tissues dynamically modulate oxidative stress caused by sepsis, and shed light on future exploration for potential diagnostics and therapeutics.

**Figure 6.**
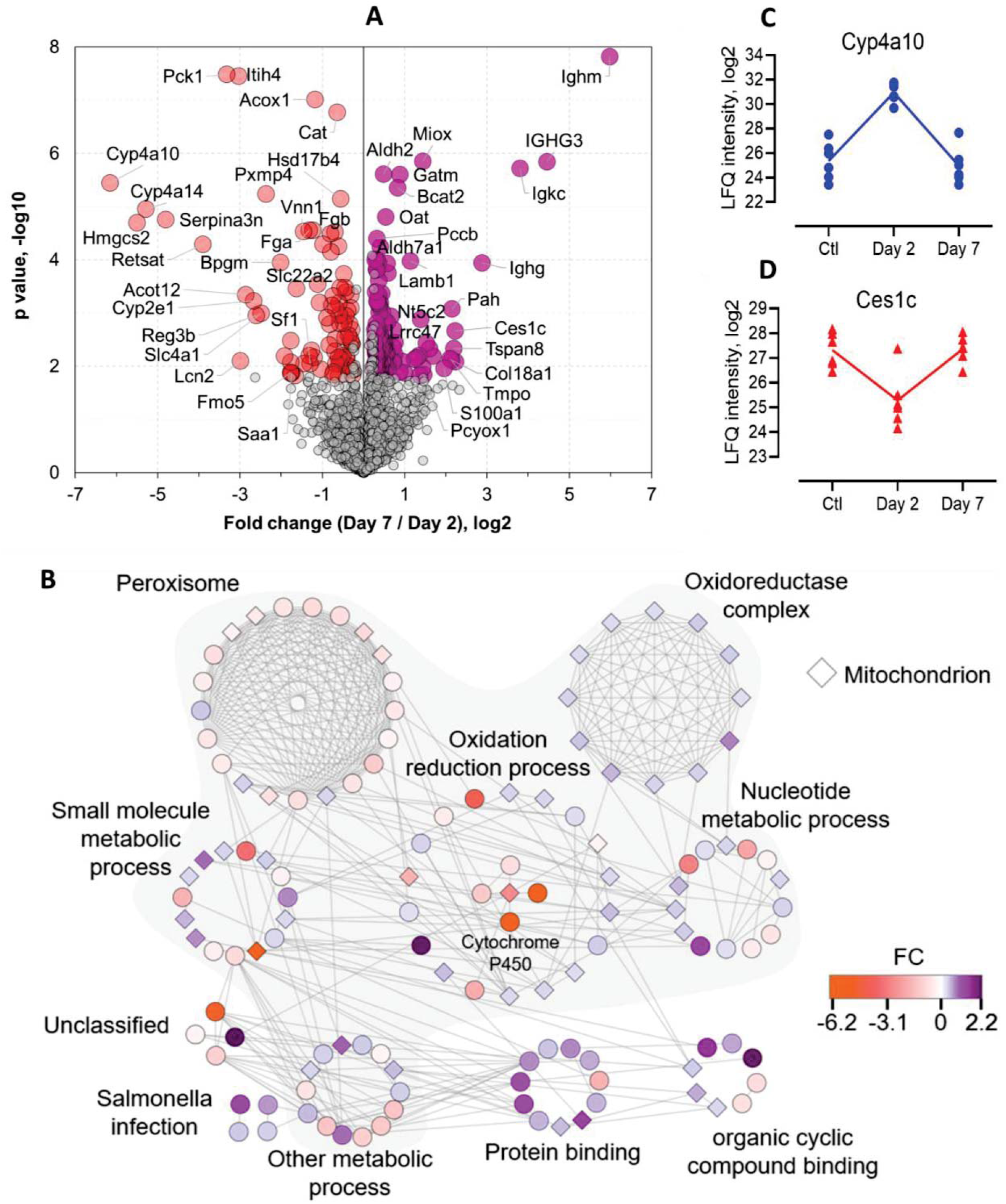
Metabolic modifications in kidneys occur upon CLP-induced sepsis. **(A)** Volcano plot shows the p value (y-axis) and fold change (x-axis) of proteins identified in both groups of kidneys. Significant proteins are highlighted in red and purple for up- and down-regulated proteins, respectively (Permutation FDR 0.05; FC ≥ 1.2). **(B)** Interaction network of the significant proteins as shown in **A**. The network and enrichment results were derived from String App in Cytoscape software. The score cutoff for interaction confidence was set to 0.9. Shaded clusters are involved in metabolic processes. The color coding is in accordance to fold changes between day 7 and day 2. Diamond shape depicts mitochondrion associated proteins. **(C-D)** Intensity plot of two representative proteins associated with the oxidation-reduction process.

## Discussion

In the present study, we have profiled the septic kidney proteome and phosphoproteome using a CLP mouse model established in our laboratory. CLP was performed in six mice (biological replicates) in each of three groups to better control reproducibility. The tissue samples were processed following an STrap approach ^89^. The STrap filters were in-house packed using bulk glass fiber membranes, which are nearly cost-free and as effective as commercial filters ^90^. The sample was analyzed in a ‘single-shot’ injection with label-free quantitation based on MaxLFQ algorithm. Our data showed minimal technical variations, thus offering critical insights into the molecular level alterations of renal proteins upon septic injury. Our CLP procedure induced only low to mid-grade severity and nearly all of the day-7 mice survived from the acute sepsis and progressed to the chronic stage. Hence, the CLP model allowed us to investigate not only the acute host response (at day 2) but also the “acute-to-chronic” transition even after moderate kidney injury, the latter of which has not been well defined molecularly ^91,92^. This transition from the acute to chronic stage reflects the refractoriness of septic kidney diseases in the clinical setting. Our study provides solid molecular evidence for the first time derived directly from septic kidneys.

S-AKI can occur in the absence of overt signs of hypoperfusion or clinical signs of hemodynamic instability. Therefore, early biomarkers are urgently needed to permit timely diagnosis and predict injury severity ^93^. However, sepsis is usually associated with multiorgan injuries, and traditional markers of S-AKI originated from plasma and urine (e.g., creatinine, urinary concentrating ability) are susceptible to interference from non-renal factors, thus are not sensitive and specific ^93^. Our global proteome and phosphoproteome data agree largely with previous urinary discoveries in the context of many known AKI markers such as NGAL. In addition, we provided a valuable source of candidates for future validation studies including Hmgcs2, Serpina3n, Cp, Chil3, Ces1c, and others. The comprehensive global analyses revealed that at the AKI stage, immune-related proteins were up-regulated while metabolism-related proteins were repressed in septic kidneys. Our data suggests that multiple immune pathways could be activated in kidneys after septic injury, including adaptive DAMPs responses, CYP-mediated lipid metabolism, peroxisome-mediated oxidative stress, and likely the inflammasome-mediated pyroptosis pathway. Unfortunately, although Gsdmd protein was found to be significantly increased at day 7 by the immunoblot assay (**Figure 1, K-L**), our untargeted MS approach was unable to detect it. Gsdmd-dependent pyroptosis cell death has been revealed for the first time in non-septic AKI in a very recent study ^43^. Our data indicated that Gsdmd expression remained at a low level at the acute stage and drastically increased at the chronic phase. Adding another time point midway through this transition (for instance, at day 4 or day 5) would likely reveal the onset of pyroptosis, which can be considered in the future studies. Nevertheless, our findings implied the potential role of pyroptosis in initiating tubular cell damage, thus deteriorating CKD, or alternatively, plasma membrane repair can be initiated to counterbalance pyroptosis (for instance, through ESCRT shedding system) ^94,95^. These questions are certainly of great interest for further investigation.

Among the responsive proteins to septic injury, Chil3 was found to be of particular interest. Chil3 (also known as YKL-40 in humans and BRP-39 in mice) does not contain known chitinase activity, though it has been implicated in host defense, inflammation, and remodeling processes ^73,74^. Chil3 has been reported as a biomarker for skin wound infection ^96^, cardiovascular disease and diabetes ^97^, and cancer ^98^. A handful of studies have shown that Chil3 has critical functional relevance to kidney ischemic injury ^99^ and renal fibrosis ^100^. There is also evidence showing its potential in predicting the onset and recovery of acute kidney injuries ^75,100,101^. However, the majority of these studies investigated chitinase secretion into the urine and used the urinary form as an indicator of disease status, which might be complicated by other non-kidney diseases such as urothelial carcinoma ^102^. Emerging studies have also started to demonstrate the mechanism of Chil3 in mediating immunity, including its role in IL-17 production in γδ T cells ^103^, and recruitment of macrophages and neutrophils in colorectal cancer ^104^. A recent study showed that Chil3 interacts with interleukin-13 receptor α2 and activates pathways that regulate apoptosis, pyroptosis, inflammasome formation, and antibacterial responses ^105^. Our data showed that Chil3 has a strong correlation with S-AKI disease onset and severity. Chil3 was elevated almost 120-fold at day 2 and more than 150-fold at day-7 after S-AKI, implying that it was involved in the progression of septic kidney injures. It remains unclear, however, whether the expression Chil3 is linked to the direct mircobial migration from gut to kidney.

Furthermore, plenty of metabolism-related proteins were significantly changed during the acute septic injury stage, which is consistent with previous findings that changes in metabolism and mitochondrial energetics occurred in response to renal damage ^106,107^. Of note, the general reduction of phosphorylation after septic injury indicates that energy usage has been reprioritized in renal tissue. The drastic increase of Hmgcs2 at day 2 suggests that ketogenesis is highly up-regulated during acute kidney injury. Similar up-regulation of Hmgc2 in diabetic kidney and heart encourages us to speculate that the energy-producing pathway in kidneys during S-AKI may be ketogenesis. Therefore, we can imagine that our CLP procedure in this study merely generated a moderate S-AKI model, and it remains intriguing to see if, for example, during a severe S-AKI senario, or at an earlier time point post-CLP, more drastic changes in these marker proteins will be seen.

In summary, our study provides the first characterization of how moderate gut injury triggered sepsis promotes S-AKI and the molecular determinant alterations *in situ*. The findings from our study will facilitate our understandings of septic kidney. The comprehensive data set will serve as a valuable resource for sepsis biomarker discovery in the future.

## Acknowledgments

We would like to acknowledge JCVI internal startup funds to Y.Y. and G.J.N.. D.Z. was supported by the National Institutes of Health (NIH) grant K01DK116816. We would like to thank Dr. Karen E. Nelson for manuscript reviewing and useful discussions.

## Data Availability

All mass spectrometry raw data associated with the study, along with detailed tables of MaxQuant output files reporting m/z, charge, identification, quantitation and phosphosite scores, and other details for all peptide identifications, have been deposited to the ProteomeXchange Consortium with the dataset identifier PXD017045 (Reviewer account details: Username; Password: pYCL0×61).

## Abbreviations

AKI: Acute kidney injury
BAD: Bcl2-associated agonist of cell death protein
CKD: Chronic kidney disease
CLP: Cecal ligation and puncture
DAMPs: Damage-associated molecular patterns
ELISA: Enzyme-linked immunosorbent assay
ESCRT: Endosomal sorting complexes required for transport
FADD: FAS-associated death domain protein
FC: Fold change
FDR: False discovery rate
GO: Gene Ontology
Gsdmd: Gasdermin D
ICU: Intensive care unit
KEGG: Kyoto Encyclopedia of Genes and Genomes
LFQ: Label-free quantitation
NGAL: Neutrophil gelatinase-associated lipocalin protein
PAMPs: Pathogen-associated molecular patterns
PAS: Periodic acid-Schiff
PCA: Principle component analysis
S-AKI: Sepsis-induced acute kidney injury
SLC: Solute carrier proteins
STrap: Suspension Trapping
TLRs: Toll-like receptors

